# The peroxisomal transporter ABCD3 plays a major role in dicarboxylic fatty acid metabolism

**DOI:** 10.1101/2021.07.26.452046

**Authors:** Pablo Ranea-Robles, Hongjie Chen, Brandon Stauffer, Chunli Yu, Dipankar Bhattacharya, Scott L. Friedman, Michelle Puchowicz, Sander M. Houten

## Abstract

Peroxisomes metabolize a specific subset of fatty acids, which include dicarboxylic fatty acids (DCAs) generated by ω-oxidation. Data obtained *in vitro* suggest that the peroxisomal transporter ABCD3 (also known as PMP70) mediates the transport of DCAs into the peroxisome, but *in vivo* evidence to support this role is lacking. In this study, we studied an *Abcd3* KO mouse model generated by CRISPR-Cas9 technology using targeted and untargeted metabolomics, histology, immunoblotting, and stable isotope tracing technology. We show that ABCD3 functions in DCA metabolism and uncover a novel role for this peroxisomal transporter in lipid metabolic homeostasis. The *Abcd3* KO mouse presents with lipodystrophy, increased circulating free fatty acids, decreased ketone bodies, enhanced hepatic cholesterol synthesis and decreased hepatic *de novo* lipogenesis. Moreover, our study suggests that DCAs are metabolized by mitochondrial β-oxidation when ABCD3 is not functional, reflecting the importance of the metabolic compartmentalization and communication between peroxisomes and mitochondria. In summary, this study provides data on the role of the peroxisomal transporter ABCD3 in hepatic lipid homeostasis and DCA metabolism, and the consequences of peroxisomal dysfunction for the liver.

## Introduction

ABCD3 (also known as 70-kDa peroxisomal membrane protein; PMP70) is one of 3 peroxisomal ATP-binding cassette (ABC) transporters (Kamijo et al. 1990). These transporters are required for the transport of substrates for peroxisomal metabolism across the peroxisomal membrane, including very long-chain fatty acids, branched-chain fatty acids like phytanic acid, the bile acid precursors di- and trihydroxycholestanoic acid (DHCA and THCA), and long-chain dicarboxylic fatty acids (DCAs) (Chornyi et al. 2021). ABCD3 deficiency (congenital bile acid synthesis defect-5; MIM 616278) is a very rare condition currently diagnosed in only one patient with severe liver disease (Ferdinandusse et al. 2015). This patient had increased plasma bile acids in particular of DHCA and THCA, which have 27 carbons and require peroxisomal β-oxidation for conversion into mature C24 bile acids. This illustrates the role of ABCD3 in transporting bile acid precursors across the peroxisomal membrane. The ABCD3 deficiency was caused by a deletion involving the last exon of the *ABCD3* gene that led to a C-terminal deletion of 24 amino acids abrogating the expression of the protein (Ferdinandusse et al. 2015).

*In vitro* studies have suggested that the CoA esters of long-chain unsaturated-, long branched-chain- and long-chain dicarboxylic- fatty acids are also preferred substrates for ABCD3 (van Roermund et al. 2014). Studies in fibroblasts from the ABCD3-deficient patient as well as phytol loading in an *Abcd3* KO mouse model, have confirmed a role for ABCD3 in the transport of branched-chain fatty acids (Ferdinandusse et al. 2015). Previous studies from our laboratory using CRISPR-Cas9 genome editing in HEK-293 cells confirmed an essential role for ABCD3 in the peroxisomal β-oxidation (FAO) of lauric, palmitic and hexadecanedioic acid (Violante et al. 2019; Ranea-Robles et al. 2021b). Even though an important role for ABCD3 in DCA metabolism has been hypothesized based on the available *in vitro* evidence, this has not yet been addressed *in vivo*.

Under specific physiological or pathological conditions such as fasting or in mitochondrial fatty acid β-oxidation (FAO) defects, long-chain DCAs are formed from long-chain fatty acids by ω-oxidation, in a process that is initiated by cytochrome P450 enzymes of the CYP4A family (PREISS and BLOCH 1964; Hardwick et al. 1987). These long-chain DCAs are specifically metabolized by peroxisomal FAO to generate acetyl-CoA and chain-shortened DCAs (Kølvraa and Gregersen 1986; Suzuki et al. 1989; Ferdinandusse et al. 2004; Dirkx et al. 2007; Ranea-Robles et al. 2021b). It has been well established that the peroxisomal enzyme enoyl-CoA hydratase/3-hydroxy acyl-CoA dehydrogenase (EHHADH; L-bifunctional protein) plays an important role in DCA β-oxidation during fasting, dietary intake of medium-chain fatty acids, and in the context of a mitochondrial FAO defect (Houten et al. 2012; Ding et al. 2013; Ranea-Robles et al. 2021b). It is generally assumed that adipic acid (C6-DCA) is a main end-product of peroxisomal DCA β-oxidation, but succinic acid (C4-DCA) is also produced (Tserng and Jin 1991; Jin et al. 2015).

Here, we have characterized an *Abcd3* KO mouse model and studied DCA metabolism using a combination of untargeted and targeted metabolomics, and stable-isotope labeling studies. We demonstrate the *in vivo* relevance of ABCD3 in the metabolism of DCAs. Altogether, our findings indicate that ABCD3 has an essential function in the transport of DCAs into the peroxisome and suggest that ABCD3 plays a previously unrecognized role in the maintenance of hepatic fatty acid and cholesterol homeostasis.

## Material and Methods

### Animal studies

Mouse experiments were approved by the Institutional Animal Care and Use Committee (IACUC) of the Icahn School of Medicine at Mount Sinai (Protocols: IACUC-2014-0061 and IACUC-2014-0100) and comply with the Guide for the Care and use of Laboratory Animals (NIH Publications No. 8023, 8^th^ edition, 2011). Mice were given free access to water and standard chow (PicoLab Rodent Diet; LabDiet, St. Louis, MO, USA) in pathogen-free housing under 12-hour light/dark cycles.

The *Abcd3* knockout (KO) mouse (Abcd3^em1(IMPC)J^) on a C57BL/6NJ background was generated by the Knockout Mouse Phenotyping Program (KOMP2) (Dickinson et al. 2016). Two male and two female heterozygous mice (*Abcd3*^+/−^) were obtained from The Jackson Laboratory and bred to produce WT and *Abcd3* KO animals. The *Abcd3* KO allele was generated at The Jackson Laboratory by injecting Cas9 RNA and 4 guide sequences GTTCATTCACTTTCTCTTGT, AACAACAATGCTTAACTACA, TTAACATTCTGCAATGCATT and CAGTAACTTGGTGAAGCTCT, which resulted in a 219 bp deletion (chr3:g.121,791,679_121,791,897del (GRCm38/mm10)) covering exon 4 and 130 bp of flanking intronic sequences including the splice acceptor and donor sites, and is predicted to cause a change of amino acid sequence after residue 82 and early stop 27 amino acids downstream. The *Abcd3* allele was genotyped using a PCR assay with the following primers: CATGTCTCATGGTGCCTGAC and ATGCTCTATGGGGCCTTTAAC, which results in a 261 bp fragment when amplifying the mutant allele, and a 483 bp fragment when amplifying the WT allele.

Five different cohorts of animals were used for the following experiments: One cohort consisted of 12 WT (6 males and 6 females, average age = 4.2 months) and 12 *Abcd3* KO (5 males and 7 females, average age = 4.2 months) mice, which were used for obtaining urine, plasma and tissue samples in the “fed”, “overnight fasted” and “photophase period fasting + L-aminocarnitine (L-AC)” conditions. The livers used for the untargeted metabolomics dataset also came from this first cohort of mice. Some of the urine samples analyzed came from mice in other cohorts. A second set of mice consisted of 5 WT (4 males and 1 female, average age = 5.3 months) and 6 *Abcd3* KO (5 males and 1 female, average age = 5.3 months) mice, used for the L-AC experiment in overnight fasted mice. A third set of mice consisted of 4 WT (2 males and 2 females, average age = 10.5 months) and 4 *Abcd3* KO (2 males and 2 females, average age = 9.7 months) mice used for the liver slices experiment. A fourth cohort of mice consisted of 12 WT (6 males and 6 females, average age = 7.1 months) and 12 *Abcd3* KO (6 males and 6 females, average age = 7.0 months) mice. This fourth cohort was used to measure cholesterol synthesis and *de novo* lipogenesis (DNL) *in vivo* using the heavy water method (^2^H_2_O) (Brunengraber et al. 2003; Bederman et al. 2009), essentially as described before (Ranea-Robles et al. 2021b). A fifth cohort of mice consisted of 5 WT (3 males and 2 females, average age = 15.9 months) and 4 *Abcd3* KO (1 males and 3 females, average age = 14.7 months) mice, used for the histologic study of the liver in older animals. EchoMRI was performed in selected mice to determine body composition measurements of fat, lean, free water, and total water masses.

For overnight food withdrawal experiments, food was removed and mice were placed in in a clean cage at 6 p.m., just before the dark cycle starts. Early the next morning, blood glucose was measured using Bayer Contour blood glucose strips. Spontaneously voided urine was collected by scruffing the mice and placing them over a plastic wrap to collect the specimen. Mice were euthanized by CO_2_ inhalation. Immediately afterwards, blood was collected from the inferior vena cava for the preparation of EDTA plasma. Organs were snap-frozen in liquid nitrogen and stored at −80°C for future analysis.

For the L-AC experiments, animals received an intraperitoneal injection of vehicle (0.9% NaCl) or L-AC (16 mg/kg) at 6 p.m., before the 12-hr dark cycle, followed by overnight food withdrawal. In a second experiment, animals received either vehicle or L-AC (16 mg/kg i.p.) at 9 a.m., followed by food withdrawal for 8 hours (photophase-period fasting). Urine, blood and organs were collected as described above.

For cholesterol synthesis and *de novo* lipogenesis (DNL) studies, mice were injected with deuterium oxide (^2^H_2_O) as 0.9% NaCl at 7 a.m., aiming to reach 4% enrichment of total body water, as previously described (Ranea-Robles et al. 2021b). Animals were given free access to chow and water during 7 hours and then a small amount of blood was drawn from the tail to measure ^2^H_2_O enrichment. Immediately after, mice were euthanized by CO_2_ inhalation. The liver was quickly frozen with a freeze-clamp, wrapped in aluminum foil, temporarily stored in liquid nitrogen, and ultimately transferred and stored at −80°C.

### Histology and immunofluorescence

Livers were collected after euthanasia and fixed by immersion in 10% formalin (Thermo Fisher Scientific) for 24 hours, washed in PBS and transferred to 70% EtOH in water until they were embedded in paraffin blocks. Serial sections (4 μm thick) were cut with a microtome. Hematoxylin & Eosin (H&E) and periodic acid Schiff (PAS) staining were performed following standard protocols, and the sections were analyzed by light microscopy. EHHADH immunofluorescence to detect hepatic peroxisomes was performed as previously described (Ranea-Robles et al. 2021a) using a specific anti-EHHADH antibody (dilution 1:50, GTX81126, Genetex). Microscopy images were taken with a Nikon Eclipse 80i microscope and the NIS-Elements BR 5.20.01 software (Nikon). Images were analyzed with ImageJ (Schindelin et al. 2012).

### Metabolite analysis / Metabolic phenotyping

Mouse plasma acylcarnitines were measured after derivatization to butylesters according to a standardized protocol (Violante et al. 2019). Tissue acylcarnitines were measured in freeze-dried tissue samples (approximately 50 mg of liver) after derivatization to propylesters essentially as described (van Vlies et al. 2005; Ranea-Robles et al. 2020). Organic acids from mouse urine were analyzed by GC-MS as previously described (Ranea-Robles et al. 2021b).

Plasma bile acids were analyzed by LC-negative ion electrospray MS/MS as previously described (Bootsma et al. 1999; Ferdinandusse et al. 2005).

Total glycerol (i.e. free glycerol and triglycerides, TR22421, Thermo Scientific), total cholesterol (TR13421, Thermo Scientific), non-esterified fatty acids (NEFA-HR(2), Wako), β-hydroxybutyrate (β-OHB, MAK-041, Sigma Aldrich), alanine aminotransferase (ALT, TR71121, Thermo Scientific) and aspartate aminotransferase (AST, TR70121, Thermo Scientific) plasma levels were measured using commercial kits following manufacturer instructions.

### Metabolic tracing of [U-^13^C]-C12-DCA in precision-cut mouse liver slices

Mouse livers were isolated from fed WT and *Abcd3* KO adult mice (2 male and 2 females per group) between 7 and 11 a.m. and precision-cut liver slices (diameter 8 mm, thickness 250 μm) were prepared as previously described (Ranea-Robles et al. 2021b). After slicing, samples were preincubated 1-2 hours inside the incubator in WEGG medium + 25 mM glucose, and randomly selected for the vehicle, [U-^13^C]-C12-DCA, or [U-^13^C]-C12-DCA + L-AC treatment. Slices were then incubated 4 hours in cell culture inserts in 6-well plates with fresh WEGG medium containing 0.4 mM carnitine and either vehicle (nothing was added), 250 μM [U-^13^C]-C12-DCA or 250 μM [U-^13^C]-C12-DCA + L-AC. WEGG medium was supplemented with 25 mM glucose in all cases. Liver slices were stored at −80°C and the medium was stored at −20°C for metabolite analysis. ^13^C-Labeled C8-DC-carnitine (M+8) and C10-DC-carnitine (M+10) were analyzed from the media and liver slice homogenates using acylcarnitine analysis as described above.

### Untargeted metabolomics

Global metabolite profiling (mView) from liver samples of 7 WT (3 males and 4 females, average age = 3.8 months) and 7 *Abcd3* KO (3 males and 4 females, average age = 3.7 months) was performed by Metabolon, Inc. (Research Triangle Park, NC), as previously described (Ranea-Robles et al. 2020, 2021a). To remove protein, to dissociate small molecules bound to protein or trapped in the precipitated protein matrix, and to recover chemically diverse metabolites, proteins were precipitated with methanol under vigorous shaking for 2 min (Glen Mills GenoGrinder 2000) followed by centrifugation. The resulting extract was analyzed by two separate reverse phase (RP)/UPLC-MS/MS methods with positive ion mode electrospray ionization (ESI), one RP/UPLC-MS/MS method with negative ion mode ESI, and one HILIC/UPLC-MS/MS method with negative ion mode ESI, as previously described (Miller et al. 2015). The scaled imputed data (Scaled Imp Data) represent the normalized raw area counts of each metabolite rescaled to set the median equal to 1. Any missing values were imputed with the minimum value. Metabolite pathway enrichment analysis using significantly altered metabolites was performed using MetaboAnalyst platform imputing KEGG IDs that were significantly different between WT and *Abcd3* KO liver (p value < 0.05) (**Table S3**) (Xia and Wishart 2011; Chong et al. 2018).

### Immunoblot

Livers were lysed in RIPA buffer supplemented with protease and phosphatase inhibitors (Thermo Fisher Scientific) using a TissueLyser II (Qiagen), followed by sonication and centrifugation (10 min at 12,000 rpm at 4°C). Protein concentration was determined by the BCA method. Proteins were separated on Bolt™ 4-12% Bis-Tris Plus gels or 7% Tris-Acetate gels (Invitrogen, Thermo Fisher Scientific), blotted onto a nitrocellulose membrane (926-31092, LI-COR) and detected as previously described (Ranea-Robles et al. 2021b). The following primary antibodies were used: anti-ABCD3 (PA1-650, Invitrogen) anti-ACOX1 (ab184032, Abcam), anti-EHHADH (GTX81126, Genetex), anti-CROT (NBP1-85501, Novus Bio), anti-CPT2 (26555-1-AP, Proteintech), anti-MCAD (55210-1-AP, Proteintech), anti-CYP4A10 (PA3-033, Invitrogen), anti-HMGCR (AMAb90618, Atlas Antibodies), anti-phospho-ACC (Ser79, 3661, Cell Signaling), anti-ACC (3662, Cell Signaling), anti-MLYCD (SAB2702043, Sigma), and anti-α-tubulin (32-2500, Thermo Fisher). The anti-MVK antibody was a gift from Dr. Hans Waterham (AMC, Amsterdam) (Hogenboom et al. 2004a, b).

### Cholesterol synthesis and de novo lipogenesis by the ^2^H_2_O method

The contribution of cholesterol synthesis (^2^H-labeled cholesterol) to the pool of total hepatic cholesterol, and the contribution of DNL (^2^H-labeled TG-bound palmitate, oleate and stearate) to the pool of total hepatic TG-bound fatty acids in mice labeled with ^2^H_2_O were determined using gas chromatography-mass spectrometry (GC-MS), as previously described (Brunengraber et al. 2003; Bederman et al. 2009; Ranea-Robles et al. 2021b). These measurements were performed at the Metabolic Phenotyping Mass Spectrometry Core (MPMS), UTHSC, Memphis, TN.

### Statistical analysis

Data are displayed as the mean ± the standard deviation (SD) with individual values shown. Differences were evaluated using unpaired t-test with Welch’s correction, or two-way ANOVA in GraphPad Prism 9, as indicated in the figure legends.

## Results

### Characterization of an Abcd3 KO mouse model

The *Abcd3* KO mouse model studied in the current work is not the same model as previously reported (Ferdinandusse et al. 2015). This *Abcd3* KO mouse model was generated and phenotyped as part of the International Mouse Phenotyping Consortium (IMPC) (Dickinson et al. 2016). There are 10 significantly affected phenotypes listed on the publicly available IMPC website including improved glucose tolerance, decreased total body fat amount, increased lean body mass and decreased bone mineral density amongst others. In our cohorts, *Abcd3* KO mice appeared healthy and developed normally. The genotype distribution in the weaned progeny (274 pups) of heterozygote breeding pairs did not follow a Mendelian distribution with a shortage of *Abcd3* KO mice (p < 0.001, **Table S1**). These data suggest that ABCD3 deficiency affects fetal and/or early postnatal development. We confirmed the absence of ABCD3 protein in the liver of adult *Abcd3* KO mice (**Fig. 1A**).

**Figure 1.**
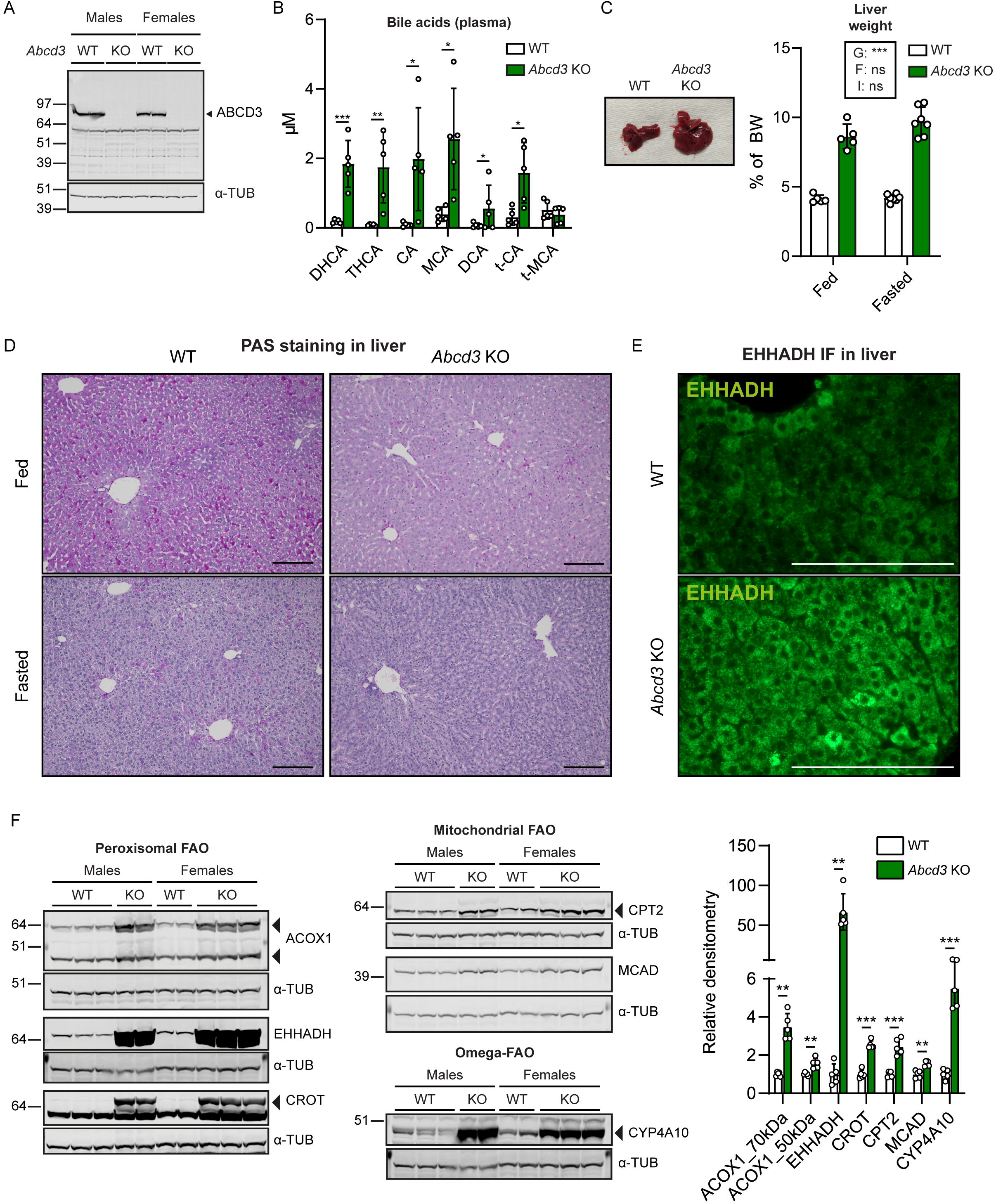
Characterization of a second *Abcd3* KO mouse model. **A**) Immunoblot using an antibody against ABCD3 in WT and *Abcd3* KO liver extracts. Alpha-tubulin (α-tub) was used as the loading control. **B**) Bile acid analysis (in μM) in plasma of WT and *Abcd3* KO fed mice (n=5). DHCA: Dihydroxycholestanoic acid; THCA: Trihydroxycholestanoic acid; CA: cholic acid; MCA: muricholic acid; DCA: deoxycholic acid; t-: taurine-. **C**) Representative images of WT and *Abcd3* KO livers (WT fed n=4, *Abcd3* KO fed n=5, WT fasted n=5, *Abcd3* KO fasted n=7) with corresponding quantification of liver-to-BW (in %). BW: body weight. The effects in a two-way ANOVA were indicated as follows: G: Genotype. F: Feeding. I: Interaction. **D**) Representative images of intracellular hepatic glycogen evaluation with PAS staining in WT and *Abcd3* KO fed and fasted livers. Scale bar = 100 μm. **E**) Representative immunofluorescent images using an antibody against EHHADH to label peroxisomes in WT and *Abcd3* KO liver sections. Scale bar = 100 μm. **F**) Immunoblot analysis using antibodies against ACOX1, EHHADH, CROT, CPT2, MCAD and CYP4A10 in liver homogenates from WT (n=5) and *Abcd3* KO (n=5) fed mice. Alpha-tubulin (α-tub) was used as loading control. EHHADH and CYP4A10 were detected on the same membrane. ACOX1 and HMGCR from Fig. 4C were detected on the same membrane. CPT2 and MVK from Fig. 4C were detected on the same membrane. MCAD and MLYCD from Fig. 4E were detected on the same membrane. Individual values, the average and the standard deviation are graphed. * p<0.05; ** p<0.01; *** p<0.001 (unpaired, two-tailed Student’s t-test in B, and F; two-way ANOVA in C).

Similar to previously reported work and consistent with an important role of ABCD3 in the transport of bile acid precursors into the peroxisome for their maturation, this *Abcd3* KO mouse model displayed pronounced elevations of the C27 bile acid precursors DHCA and THCA in plasma (**Fig. 1B**). The primary bile acids cholic acid (CA) and muricholic acid (MCA) were increased in plasma (**Fig. 1B** and **Table S2**) consistent with cholestasis as was reported in the ABCD3-deficient patient (Ferdinandusse et al. 2015). We noticed a pronounced hepatomegaly in *Abcd3* KO mice with no histological signs of increased fat or fluid (**Fig. 1C, S1A**), similar to previous findings (Ferdinandusse et al. 2015). The plasma levels of the hepatic enzymes ALT and AST tended to be slightly elevated in *Abcd3* KO mice (**Fig. S1B**), which may indicate mild liver injury in *Abcd3* KO mice. We noted hepatic nodules with vesicular steatosis and inflammatory infiltrates in 2 out of 4 15-month-old *Abcd3* KO mice (**Fig. S1C**). None of the 5 WT mice analyzed had these nodules. Notably, PAS staining uncovered a marked glycogen deficiency in *Abcd3* KO fed mice (**Fig. 1D**). The difference in PAS staining was less pronounced upon overnight fasting likely due to depletion of glycogen stores in WT livers (**Fig. 1D**). We observed a reduction in total glycogen synthase (GYS) protein levels in fed *Abcd3* KO mice with no changes in the inhibitory phosphorylation of GYS on residue Ser641 (**Fig. S1D**). These findings suggest decreased glycogen synthesis and/or increased glycogen utilization in *Abcd3* KO livers.

Despite the changes in liver glycogen, we did not observe any changes in blood glucose levels between WT and *Abcd3* KO mice in the fed or in the fasted state (**Fig. S1E**). Plasma non-esterified fatty acids levels were higher in fasted *Abcd3* KO mice compared with WT mice (**Fig. S1F**), whereas total glycerol was decreased in fed and fasted *Abcd3* KO mice (**Fig. S1G**), suggesting that ABCD3 deficiency leads to increased lipolysis or impaired fatty acid uptake. The increased lipolysis was further supported by a reduced amount of epididymal fat in *Abcd3* KO mice (**Fig. S1H**). Indeed, the characterization at the IMPC and our work showed decreased total body fat amount and increased lean body mass (**Fig. S1I, S1J**), suggesting that ABCD3-deficient mice have a lipodystrophic phenotype. Total cholesterol levels in plasma were reduced in fed *Abcd3* KO mice, but were increased in fasted *Abcd3* KO as compared with WT mice in the same state (**Fig. S1K**). The plasma levels of the ketone body β-hydroxybutyrate (β-OHB) did not increase in *Abcd3* KO mice after fasting, in contrast to the expected fasting-induced increase in β-OHB we observed in WT mice (**Fig. S1L**), which suggests impaired ketogenesis or deficient ketolysis.

### Upregulation of fatty acid oxidation pathways in Abcd3 KO mouse liver

The increase in liver size in *Abcd3* KO mice may be caused by peroxisome proliferation (Reddy and Krishnakantha 1975), which is typically caused by the induction of peroxisomal, as well as mitochondrial and ER enzymes involved in FAO. EHHADH immunostaining revealed increased peroxisomal density in *Abcd3* KO livers compared with WT livers (**Fig. 1E**). Indeed, we found a pronounced induction of the peroxisomal FAO proteins ACOX1, EHHADH, and CROT in the liver of fed *Abcd3* KO mice (**Fig. 1F**). The mitochondrial FAO pathway was increased alongside as protein levels of CPT2 and MCAD were higher in *Abcd3* KO livers (**Fig. 1F**). Fatty acid ω-oxidation was also induced in *Abcd3* KO livers, as evidenced by increased hepatic CYP4A10 protein levels (**Fig. 1F**). In summary, these data imply that ABCD3 deficiency results in hepatic peroxisomal proliferation and the induction of mitochondrial and peroxisomal β-oxidation as well as ω-oxidation pathways.

### Untargeted metabolomic analysis of WT and Abcd3 KO mouse livers

To characterize the changes in liver metabolites caused by the loss-of-ABCD3 function, we performed untargeted metabolomics in liver from overnight fasted male and female WT and *Abcd3* KO mice (n=7 per group). We identified a total of 877 named biochemicals (**Table S3**). WT and *Abcd3* KO livers were completely separated by their first principal component showing that most of the variation in the metabolome is due to the *Abcd3* KO (**Fig. 2A**). A total of 481 metabolites were significantly different (p value < 0.05) between WT and *Abcd3* KO livers, 159 of them were increased, whereas 322 of them were decreased in *Abcd3* KO livers (**Fig. 2B**).

**Figure 2.**
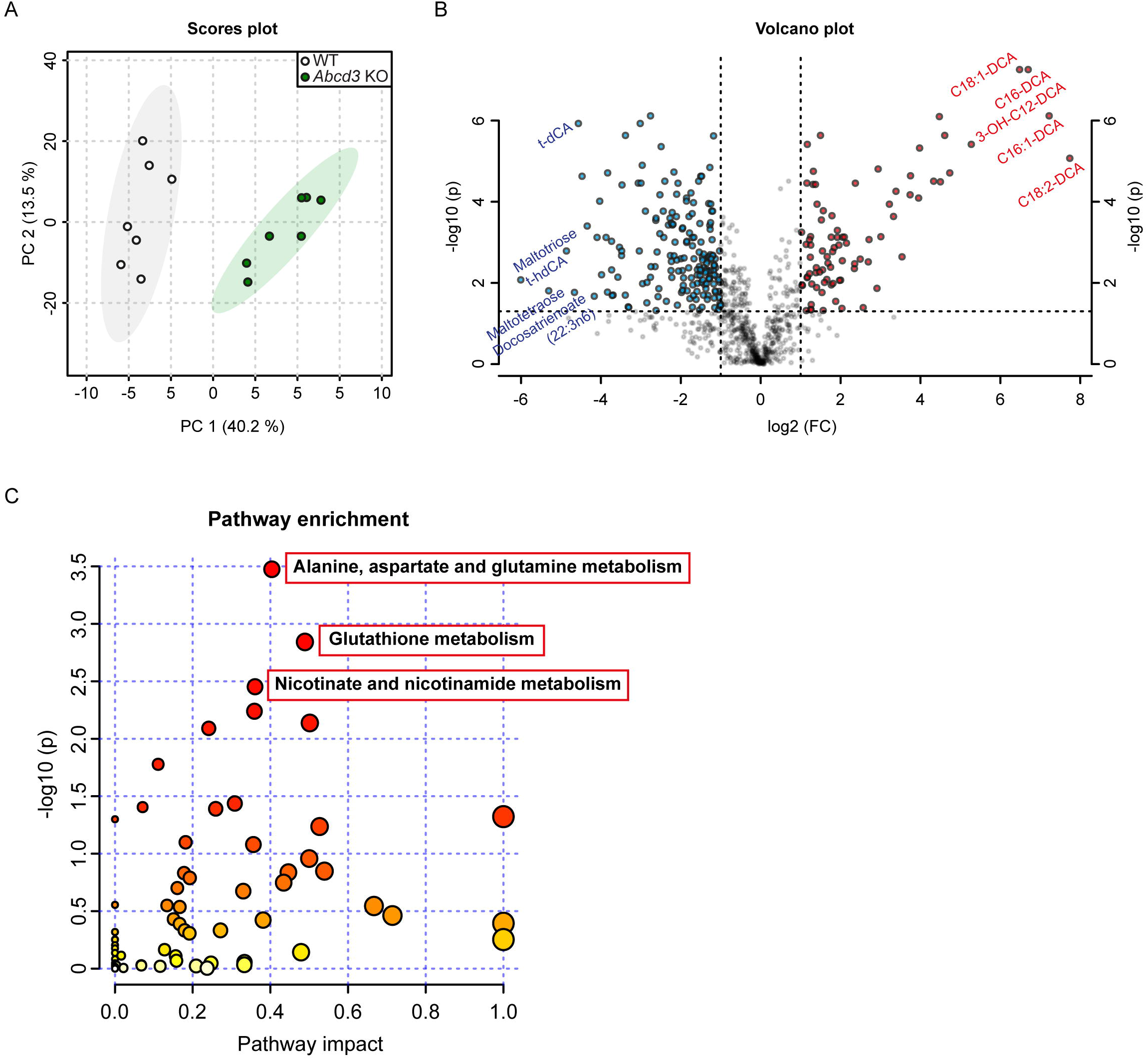
Untargeted metabolomic analysis of WT and *Abcd3* KO mouse livers. **A**) 2-D PCA scores plot between select PCs obtained from WT and *Abcd3* KO hepatic metabolome (n=7). The explained variances are shown in brackets. **B**) Important metabolites selected by volcano plot with fold-change threshold (x-axis) = 2 and p-value threshold (y-axis) = 0.05. Blue and red circles represent metabolites above the p-value threshold, and below (blue) or above (red) the fold change threshold, respectively. Both fold-changes and p-values are log transformed. The top 5 metabolites below and above the fold-change threshold are indicated. **C**) Summary of pathway enrichment analysis using significantly altered metabolites with a KEGG ID. Scatterplot represents unadjusted p-values from integrated enrichment analysis and impact values from pathway topology analysis. The node color is based on the p-values and the node radius represents the pathway impact values. The 3 significantly altered KEGG pathways using Fisher’s exact t-test (adj. p < 0.1) are indicated.

Consistent with a role of ABCD3 in DCA transport across the peroxisomal membrane (van Roermund et al. 2014; Ranea-Robles et al. 2021b), 12 DCAs were among the top-15 increased metabolites by fold-change in *Abcd3* KO livers, including long-chain DCAs (C14 to C18), with some of them displaying an increase of more than 100-fold (C18:2-DCA, C16:1-DCA, and C16-DCA) (**Fig. 2B** and **Table S3**). Primary bile acids were found among the most decreased metabolites by fold-change in *Abcd3* KO livers (**Fig. 2B** and **Table S3**), in line with the defect in bile acid biosynthesis caused by ABCD3 deficiency. Glycogen-derived metabolites, such as maltose, maltotriose, and maltotetraose, were also among the most decreased metabolites in *Abcd3* KO livers (Fig. 2B and Table S3), consistent with the glycogen depletion observed in liver sections by PAS staining (**Fig. 1G**). We also found an increase in pipecolate levels in *Abcd3* KO livers, which may be indicative of peroxisomal dysfunction (Mihalik et al. 1989) (**Table S3**). We observed a robust decrease in sphingomyelins with diverse acyl chains (20 out of the 27 detected sphingomyelins were significantly decreased in *Abcd3* KO livers) (**Table S3**). Thus, untargeted metabolomics confirmed that the loss of ABCD3 function causes a defect in the biosynthesis of primary bile acids (Ferdinandusse et al. 2015), depletes hepatic glycogen stores, and suggest that ABCD3 has an important role mediating the transport of DCAs into the peroxisome in the liver.

To determine the specific pathways in which significantly changed metabolites were involved, we performed KEGG pathway enrichment analysis. Pathways involving “Alanine, aspartate and glutamate metabolism”, “Glutathione metabolism”, and “Nicotinate and nicotinamide metabolism” were identified as the significantly enriched pathways among the altered metabolites (**Fig. 2C and Table S3**). In fact, we found a reduced availability of amino acids in *Abcd3* KO livers, with decreased levels of glycine, serine, threonine, alanine, asparagine, glutamate, tyrosine, tryptophan, and proline (**Table S3**). The term “TCA cycle” was also enriched, with a general decrease of TCA cycle intermediates such as citrate, aconitate, α-ketoglutarate, succinate, fumarate, and malate in *Abcd3* KO livers (**Table S3**). In summary, ABCD3 deficiency has a profound impact on the metabolic profile of mouse liver, characterized by an increase in DCAs, decrease in primary bile acids, and reduced amino acid availability.

### Dicarboxylic acid metabolism in Abcd3 KO mice

To further establish the essential role of ABCD3 in peroxisomal DCA metabolism, we measured urinary excretion of DCAs in an independent cohort of male and female *Abcd3* KO mice in the *ad libitum* fed state or after one night of food withdrawal (**Table S4A**). The urine of *Abcd3* KO mice contained elevated amounts of even-chain DCAs, namely adipic acid (C6-DCA), suberic acid (C8-DCA), sebacic acid (C10-DCA), and dodecanedioic acid (C12-DCA) in both fed and fasted state (**Fig. 3A, S2A, and Table S4A**). Tetradecanedioic acid (C14-DCA) was only elevated in fasted *Abcd3* KO mice (**Table S4A**). Odd-chain DCAs pimelic acid (C7-DCA) and azelaic acid (C9-DCA) were also elevated in the urine of *Abcd3* KO mice in both conditions (**Table S4A**). The excretion of C8-DCA was particularly elevated in *Abcd3* KO mice, reaching values of up to 6 mol/mol creatinine in urine of fasted male *Abcd3* KO mice (**Fig. 3A, S2A, and Table S4A**). The 3-OH forms of C8-DCA, C10-DCA, and C12-DCA were also increased in the urine of fed and fasted *Abcd3* KO mice (**Fig. S2B, S2C, and Table S4A**). 3-OH-C6-DCA levels were decreased in the urine of fasted female *Abcd3* KO mice (**Fig. S2C, Table S4A**). These results demonstrate that *Abcd3* KO mice present with a pronounced medium-chain dicarboxylic aciduria that is exacerbated upon fasting.

**Figure 3.**
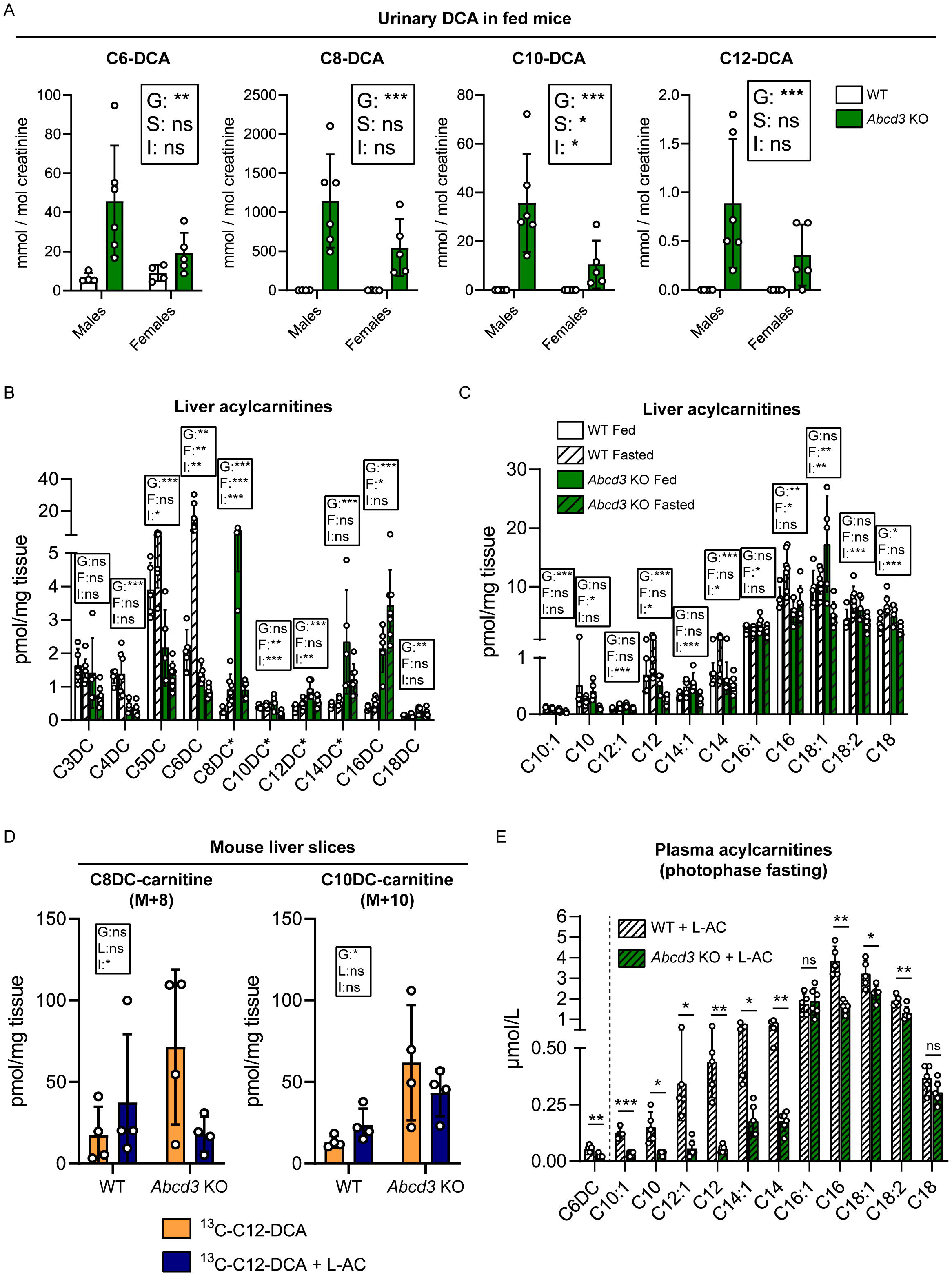
Changes in fatty acid metabolism in *Abcd3* KO mice. **A**) Urinary C6-DCA, C8-DCA, C10-DCA, and C12-DCA (in mmol/mol creatinine) in WT (n=4 males, n=4 females) and *Abcd3* KO (n=6 males, n=5 females) fed mice. **B**) Liver dicarboxylylcarnitine profile (in pmol/mg tissue) in WT (n=5 fed, n=7 fasted) and *Abcd3* KO (n=5 fed, n=7 fasted) mice. Dicarboxylylcarnitine species that could not be distinguished from other hydroxyacylcarnitine species with the same nominal mass (isobaric compounds) are marked with an asterisk, as indicated here: C8DC*/C12-OH; C10DC*/C14-OH; C12DC*/C16-OH; C14DC*/C18-OH. **C**) Liver medium- and long-chain acylcarnitine profile (in pmol/mg tissue) in WT (n=5 fed, n=7 fasted) and *Abcd3* KO (n=5 fed, n=7 fasted) mice. **D**) Measured [U-^13^C]-labeled C8-DC-carnitine, and C10-DC-carnitine (in pmol/mg of tissue) in mouse liver slices after 4-hr incubation of WT and *Abcd3* KO mouse liver slices (n=4) with [U-^13^C]-C12-DCA alone or with [U-^13^C]-C12-DCA + L-aminocarnitine (L-AC). **E**) Plasma medium- and long-chain acylcarnitine profile (in μmol/L) in WT (n=5) and *Abcd3* (n=6) KO mice treated with L-AC and subjected to food withdrawal during the photophase. Individual values, the average and the standard deviation are graphed.* p<0.05; ** p<0.01; *** p<0.001 (two-way ANOVA in A-D; unpaired, two-tailed student t-test in E). The effects in the two-way ANOVA are indicated as follows; G: Genotype; S: Sex; F: Feeding; L: L-AC; I: Interaction.

### Acylcarnitine metabolism in Abcd3 KO mice

We analyzed the plasma and hepatic acylcarnitine profile in WT and *Abcd3* KO mice, in the fed state and after overnight food withdrawal (**Table S4B, S4C**). In the liver, we found that most of the even-chain dicarboxylylcarnitines with a carbon chain of 8 carbons and longer were increased in *Abcd3* KO mice (**Fig. 3B**). C8DC-carnitine hepatic levels showed genotype- and feeding-specific effects, as their levels were only increased in the fed state when comparing *Abcd3* KO livers to WT livers (**Fig. 3B**). On the other hand, short-chain dicarboxylylcarnitines C4DC-, C5DC-, and C6DC-carnitine were lower in *Abcd3* KO liver (**Fig. 3B**). Dicarboxylylcarnitines in plasma showed a similar profile, characterized by a decrease of the short-chain dicarboxylylcarnitines C3DC-, C4DC-, C5DC-, and C6DC-carnitine, and an increase of C8DC-carnitine in *Abcd3* KO plasma (**Fig. S3A**). These results are consistent with a deficient DCA β-oxidation in *Abcd3* KO mice.

Plasma and hepatic acylcarnitine profile also revealed an effect of ABCD3 deficiency on monocarboxylylcarnitines. Levels of acetylcarnitine (C2-carnitine) and hydroxybutyrylcarnitine (C4OH-carnitine), the latter of which reflects D-3-hydroxybutyrate (Soeters et al. 2012), were reduced in plasma and liver of fasted *Abcd3* KO mice (**Fig. S3B-S3E**), suggesting a defect in the generation of acetyl-CoA and ketone bodies in the *Abcd3* KO liver. Moreover, the fasting-induced increase in hepatic free carnitine levels was prevented by the loss of ABCD3 function (**Fig. S3C**), whereas in plasma, free carnitine levels were lower in fed and fasted *Abcd3* KO mice (**Fig. S3B**). Short-chain acylcarnitine (C3-, C4-, C5-, and C6-) levels were decreased in plasma and liver of fed and fasted *Abcd3* KO mice (Fig. **S3D, S3E**). Medium- and long-chain acylcarnitines such as C12-, C14:1-, C14-, C16-, C18:1-, C18:2- and C18-carnitine were also decreased in the liver and plasma of *Abcd3* KO mice, particularly in the fasted state (**Fig. 3C, S3F**). These findings suggest that ABCD3 deficiency also impacts on other metabolic pathway such as β-oxidation of monocarboxylic acids.

### DCAs are metabolized in the mitochondria upon ABCD3 deficiency

To further investigate the defect in DCA metabolism observed in *Abcd3* KO mice, we probed DCA metabolism with [U-^13^C]-C12-DCA in precision-cut liver slices (PCLS) (Ranea-Robles et al. 2021b). We measured acylcarnitines in the PCLS tissue and media. We detected fully labeled C10DC-carnitine (M+10) and C8DC-carnitine (M+8) in WT PCLS after incubation with the C12-DCA tracer, which confirmed that C12-DCA underwent at least 2 cycles of β-oxidation in the mouse PCLS (**Fig. 3D**). The amount of fully labeled C10DC-carnitine (M+10) and C8DC-carnitine (M+8) increased in *Abcd3* KO PCLS tissue compared with WT PCLS (**Fig. 3D**). Similarly, C10DC-carnitine (M+10) was increased in *Abcd3* KO PCLS media (**Fig. S3G**). Thus similar to the *in vivo* data, the PCLS from *Abcd3* KO liver accumulate more medium-chain DCAs when compared to WT liver.

Since the accumulation of medium-chain DCAs in *Abcd3* KO livers and PCLS suggest that mitochondria metabolize DCAs upon ABCD3 loss-of-function, we incubated the PCLS with [U-^13^C]-C12-DCA and the CPT2 inhibitor L-aminocarnitine (L-AC) (Chegary et al. 2008). Under these conditions, the increase in C8DC-carnitine was partially prevented in *Abcd3* KO PCLS (**Fig. 3D**). Similarly, L-AC decreased the accumulation of C10DC-carnitine in the media (**Fig. S3G**). Together these results suggest that CPT2-mediated mitochondrial β-oxidation contributes to the accumulation of medium-chain DCAs upon ABCD3 deficiency.

To further study the potential contribution of mitochondria to DCA β-oxidation in *Abcd3* KO mice, we measured urinary DCAs and the acylcarnitine profile in WT and *Abcd3* KO mice that treated with L-AC (**Table S4A, S4D**). In a first experiment in which the mice were subjected to L-AC treatment combined with overnight food withdrawal, 4 out of the 6 *Abcd3* KO mice in the L-AC group died or had to be euthanized, whereas WT mice did not develop any complications from L-AC treatment. This indicates that a combined defect in mitochondrial and peroxisomal FAO is often fatal during fasting in mice. In urine samples collected from 5 WT and 3 *Abcd3* KO mice, we did not observe any major effects of CPT2 inhibition on the levels of C8-DCA, C10-DCA, C12-DCA, their 3-OH forms, and C14-DCA (**Fig. S3H, S3I**). In contrast, urinary C6-DCA levels decreased, whereas 3-OH-C6-DCA was not detectable in *Abcd3* KO mice upon treatment with L-AC (**Fig. S3H, S3I**). In a second experiment, WT and *Abcd3* KO mice were treated with L-AC and subjected to a maximum of 8 hours food withdrawal during the photophase, a period in which mice usually do not feed. Plasma was collected to analyze the acylcarnitine profile (**Table S4D**). *Abcd3* KO mice treated with L-AC in this group showed lower levels of medium- and long-chain saturated and mono-unsaturated acylcarnitines in plasma, compared with treated WT mice (**Fig. 3E**). Similarly to what was observed in the urine, the levels of C6DC-carnitine in plasma decreased in *Abcd3* KO mice treated with L-AC (**Fig. 3E**). Overall, these results confirm the important role of ABCD3 in the generation of medium- and long-chain acylcarnitines when mitochondrial FAO is blocked or saturated (Violante et al. 2019). At the same time, these data indicate that DCAs can be metabolized in the mitochondria when the peroxisomal transporter ABCD3 is not functional, reflecting the importance of compartmentalization for proper fatty acid metabolic homeostasis.

### ABCD3 deficiency alters hepatic cholesterol synthesis and de novo lipogenesis

To better characterize the alterations in cholesterol and lipid metabolism in *Abcd3* KO mice, we measured hepatic cholesterol synthesis and *de novo* lipogenesis (DNL) *in vivo* in fed mice using the deuterated water (^2^H_2_O) method. We observed an induction of cholesterol synthesis in male and female *Abcd3* KO mice (**Fig. 4A**) with no changes in the total hepatic cholesterol levels (**Fig. 4B**). The increased rate of cholesterol synthesis in *Abcd3* KO livers was accompanied by an induction of the cholesterol synthesis pathway, reflected by the increased levels of 3-hydroxy-3-methylglutaryl coenzyme A reductase (HMGCR) and mevalonate kinase (MVK) proteins (**Fig. 4C**). We also observed a pronounced reduction in hepatic DNL for all the TG-bound fatty acids measured (palmitate, oleate, and stearate) (**Fig. 4D**), whereas only the total levels of TG-bound stearate were lower in fed *Abcd3* KO livers (**Fig. 4E**). Phosphorylated acetyl-CoA carboxylase 1 (ACC1) on residue Ser79 (lower band) and total ACC1 levels (lower band) were increased in *Abcd3* KO mice, whereas the ratio pACC1/ACC1 was decreased (**Fig. 4F**), suggesting that AMPK-induced phosphorylation of ACC1 is not responsible for the induction of hepatic mitochondrial FAO proteins and inhibition of DNL. Malonyl-CoA decarboxylase (MLYCD) protein levels were increased in fed *Abcd3* KO livers (**Fig. 4F**). MLYCD is an enzyme that catalyzes the decarboxylation of malonyl-CoA to acetyl-CoA and thus stimulates mitochondrial FAO while inhibiting DNL (An et al. 2004). This result is consistent with the reduced levels of malonate and malonylcarnitine in the liver metabolomics and the acylcarnitine profile (**Table S1 and Table S4**). In summary, loss-of-ABCD3 function in mice causes an increase in hepatic cholesterol synthesis and a reduction in DNL, accompanied by changes in the enzymes that modulate these metabolic pathways.

**Figure 4.**
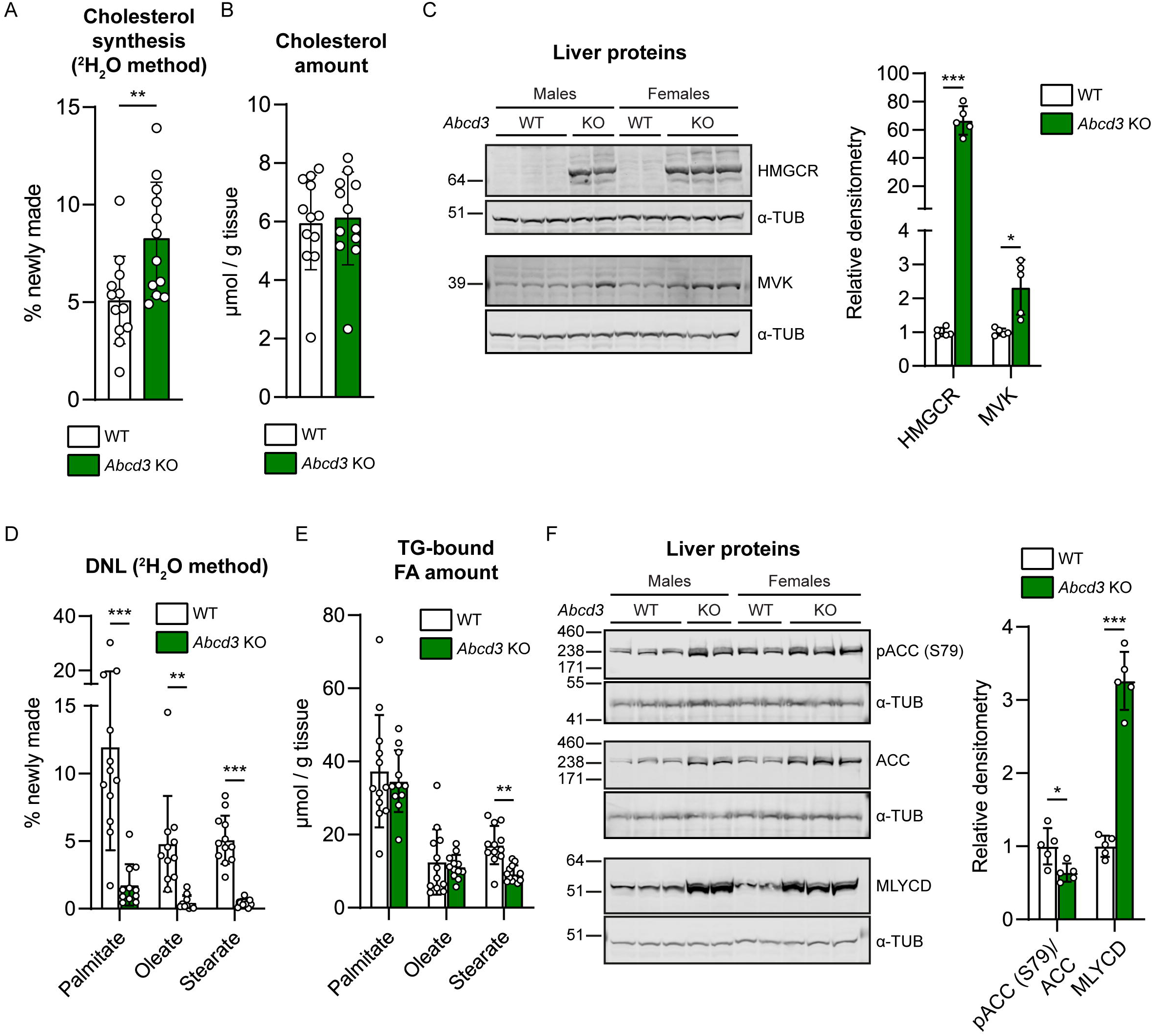
ABCD3 deficiency alters hepatic cholesterol synthesis and *de novo* lipogenesis. **A**) Cholesterol synthesis (in “% of newly made”) was calculated from the incorporation of deuterium isotopes (^2^H) into cholesterol in the liver of WT (n=6 males and n=6 females) and *Abcd3* KO (n=6 males and n=6 females) mice. **B**) Total cholesterol amount (in “μmol/g tissue”) in the liver of WT (n=6 males and n=6 females) and *Abcd3* KO (n=6 males and n=6 females) mice was measured by mass spectrometry. **C**) Immunoblot using antibodies against HMGCR and MVK in liver homogenates from WT (n=5) and *Abcd3* KO (n=5) fed mice. Alpha-tubulin (α-tub) was used as loading control. **D**) *De novo* lipogenesis (DNL, in “% of newly made”) was calculated from the incorporation of deuterium isotopes (^2^H) into the corresponding TG-bound fatty acids (palmitate, oleate, and stearate) in the liver of WT (n=6 males and n=6 females) and *Abcd3* KO (n=6 males and n=6 females) mice. **E**) TG-bound fatty acid (palmitate, oleate, and stearate) content (in “μmol/g tissue”) was measured by mass spectrometry. **F**) Immunoblot using antibodies against phosphorylated ACC on the residue Ser79 [pACC (S79)] and MLYCD in liver homogenates from WT (n=5) and *Abcd3* KO (n=5) fed mice. Total ACC or alpha-tubulin (α-tub) were used as loading controls as indicated in the x-axis of the graph. Individual values, the average and the standard deviation are graphed.* p<0.05; ** p<0.01; *** p<0.001 (unpaired, two-tailed student t-test).

## Discussion

We have characterized a second *Abcd3* KO mouse model and focused on DCA metabolism. Our studies revealed an important role of the peroxisomal transporter ABCD3 in the transport of DCAs into the peroxisome and confirmed its role in bile acid biosynthesis. In addition, we found that *Abcd3* KO mice a significant perturbation in lipid homeostasis reflected by lipodystrophy, increased circulating free fatty acids, enhanced hepatic cholesterol synthesis, decreased hepatic DNL and defective ketone body synthesis.

A role for ABCD3 protein in peroxisomal DCA import has been proposed based on *in vitro* findings (van Roermund et al. 2014; Violante et al. 2019; Ranea-Robles et al. 2021b), but has been never documented *in vivo*. The hepatic metabolome of *Abcd3* KO livers showed a pronounced increase in long-chain DCAs and long-chain dicarboxylylcarnitines. Remarkably, this was accompanied by a marked medium-chain dicarboxylic aciduria in particular of C8-DCA. The difference in DCA profiles between liver and urine suggests the involvement of a transport mechanism in liver and/or kidney that is selective for medium-chain DCA and prevents the excretion of long-chain DCAs. Overall, our data provide *in vivo* evidence to a role of ABCD3 in peroxisomal metabolism of DCAs.

Interestingly, the accumulation of medium-chain DCAs such as suberic acid (C8-DCA) in the liver and the urine of *Abcd3* KO mice suggests that DCA β-oxidation is able to proceed and is seemingly at odds with an essential role for ABCD3 in the transport of long-chain DCAs into the peroxisome for their subsequent β-oxidation. There are two possible explanations for this observation. First, the increased C8-DCA production could be of peroxisomal origin, because the two other peroxisomal ABC transporters, ABCD1 and ABCD2, may take over the function of ABCD3 and transport the DCAs into the peroxisome for β-oxidation. Although we cannot rule out such a compensatory role for ABCD1 and/or ABCD2, the low expression of these proteins in liver and kidney (Kemp et al. 2011) in combination with their decreased preference for DCAs (van Roermund et al. 2014) makes this less likely. In addition, compensation by ABCD1 and/or 2 would not explain the increased formation of medium-chain DCAs. Therefore, we hypothesize that the C8-DCA is produced by the mitochondrial FAO machinery and envision two different mechanisms. Incomplete mitochondrial FAO may lead to the release of FAO intermediates such as C8-carnitine or octanoic acid, which are ultimately converted into C8-DCA by ω-oxidation. Subsequent peroxisomal β-oxidation is defective in *Abcd3* KO mice leading to C8-DCA excretion. Alternatively, the mitochondrial FAO system may be forced to metabolize long-chain DCAs because of their accumulation due to the peroxisomal import defect. It is known that the mitochondrial FAO machinery is not efficient in handling DCAs (Pettersen 1973; Pourfarzam and Bartlett 1991, 1993; Ranea-Robles et al. 2021b), and this may be reflected in the accumulation of (3-OH) medium-chain DCA intermediates in *Abcd3* KO mice. The data obtained in the PCLS experiment support that mitochondrial β-oxidation contributes to the accumulation of medium-chain DCAs in *Abcd3* KO mice.

Our studies indicate subtle liver injury caused by ABCD3 deficiency characterized by hepatomegaly, cholestasis, slightly elevated transaminases and depletion of glycogen stores. Liver pathologies are common in peroxisome biogenesis disorders and peroxisomal β-oxidation defects, and also occur in mouse models of these diseases (Baes and Van Veldhoven 2016). Mice with non-functional peroxisomes in the liver (liver-specific *Pex5* KO mice) share many hepatic features with the *Abcd3* KO mice such as hepatomegaly, induction of PPARα signaling, impaired DNL, reduction in hepatic glycogen levels, and frequent development of liver tumors (Dirkx et al. 2005; Peeters et al. 2011b, a). We show here that the bile acid biosynthesis defect and the accumulation of medium- and long-DCAs, including odd-chain DCAs are the most striking metabolic alterations that occur in the *Abcd3* KO liver. A similar accumulation of DCAs is characteristic for peroxisomal disorders such as Zellweger spectrum disorders (ZSD), and pseudoneonatal adrenoleukodystrophy (caused by *ACOX1* mutations) (Rocchiccioli et al. 1986; Rizzo et al. 2003), which also display liver pathologies (Ferdinandusse et al. 2007; Klouwer et al. 2015). Ultimately, the redirection of DCAs to the mitochondria may negatively affect mitochondrial function as has been suggested by others (Passi et al. 1984; Tonsgard and Getz 1985). Thus, it is tempting to speculate that the accumulation of long-chain DCAs may be harmful to mitochondria, and contributes to the mitochondrial abnormalities observed in peroxisomal disorders.

The metabolome of *Abcd3* KO liver also revealed a generalized decrease in sphingomyelins, which has been recently proposed as a specific metabolic signature of ZSD (Wangler et al. 2018). Moreover, inactivation of peroxisomal enzymes involved in DCA β-oxidation, such as ACOX1 and EHHADH, causes liver disease characterized by hepatosteatosis, liver tumors, and liver fibrosis, on a standard chow diet in the case of *Acox1* KO mice (Fan et al. 1996), or triggered by a lauric acid-enriched diet in the case of *Ehhadh* KO mice (Ding et al. 2013). Altogether, these data suggest that DCA β-oxidation, in which ABCD3 plays an essential role, may be crucial for the maintenance of liver functions.

In the absence of ABCD3, the overload of metabolites transported by ABCD3 into the peroxisomes may trigger the activation of the transcription factor PPARα (Zomer et al. 2003). Indeed, we found an induction of peroxisomal and mitochondrial β-oxidation, and ω-oxidation proteins in *Abcd3* KO livers, which is consistent with an activation of PPARα. The induction of PPARα has been also reported in mice with inactivation of different peroxisomal β-oxidation enzymes, ACOX1 (Fan et al. 1996), HSD17B4 (peroxisomal D-bifunctional protein) (Martens et al. 2008), EHHADH (Ding et al. 2013) and SCPx (Seedorf et al. 1998). Since this study was performed in mice on a standard diet, the accumulation of branched-chain fatty acids is not likely to contribute to PPARα activation. Therefore, we hypothesize that DCAs are the most likely endogenous activators of PPARα in *Abcd3* KO mice (Ding et al. 2013). Therefore, this *Abcd3* KO mouse represents an excellent model to investigate the potential activation of PPARα by DCAs.

We also observed an induction of proteins involved in cholesterol synthesis and an increase in cholesterol synthesis rates in *Abcd3* KO livers. Perturbations in cholesterol homeostasis have been reported in other mouse models for peroxisomal disease such as the *Ehhadh* KO mice (Ranea-Robles et al. 2021b). Indeed it is known that acetyl-CoA produced by peroxisomal β-oxidation can be used in the cytosol to initiate cholesterol synthesis (Leighton et al. 1989; Kasumov et al. 2005). We hypothesize that the disruption in acetyl-CoA production by peroxisomal β-oxidation in the *Abcd3* and *Ehhadh* KO mouse models decreases the availability of precursors for cholesterol synthesis necessitating the increase in cholesterol biosynthetic capacity. Combined, these findings support the notion that peroxisomal DCA β-oxidation modulates hepatic cholesterol homeostasis.

In summary, *Abcd3* KO mice represent an interesting animal model to investigate the physiological role of peroxisomal DCA metabolism. In addition, our findings can be useful for the study of the pathophysiology and the diagnosis of ABCD3 deficiency in humans. Finally, this animal model provides information on the role of the peroxisomal transporter ABCD3 in hepatic lipid homeostasis and the consequences of peroxisomal dysfunction on liver pathology.

## Acknowledgements

We thank Dr. Hans R. Waterham for providing the MVK antibodies, and Dr. Frédéric M. Vaz for the bile acid analysis. We acknowledge the help of the shared resource facilities at the Icahn School of Medicine at Mount Sinai (Colony Management, the Genomics Core, the Comparative Pathology Laboratory the Biorepository and Pathology Core). Research reported in this publication was supported by the National Institute of Diabetes and Digestive and Kidney Diseases of the National Institutes of Health under Award Number R01 DK113172 (to SMH) and R01 DK128289 (to SLF). The content is solely the responsibility of the authors and does not necessarily represent the official views of the National Institutes of Health.

