## Supplementary Material for "The peroxisomal transporter ABCD3 plays a major role in dicarboxylic fatty acid metabolism"

**Supplemental information Table of Contents**

**<u>Supplementary Figure legend</u>**

**Supplementary Figure S1.** Characterization of a second *Abcd3* KO mouse

**Supplementary Figure S2.** Urinary DCA alterations in *Abcd3* KO mice

**Supplementary Figure S3.** DCA metabolism alterations in *Abcd3* KO mice

**<u>Supplementary Tables</u>**

**Supplementary Table S1.** Genotype distribution in a cross of *Abcd3*$^{+/-}$ x *Abcd3*$^{+/-}$ mice

**Supplementary Table S2.** Plasma bile acid concentrations in *Abcd3* KO fed mice

**Supplementary Table S3.** Untargeted metabolomics dataset comparing WT and *Abcd3* KO liver

**Supplementary Table S4.** Urinary DCA, and plasma and hepatic acylcarnitine concentrations in WT and *Abcd3* KO mice.

### Supplementary Figures

#### Supplementary Figure 1

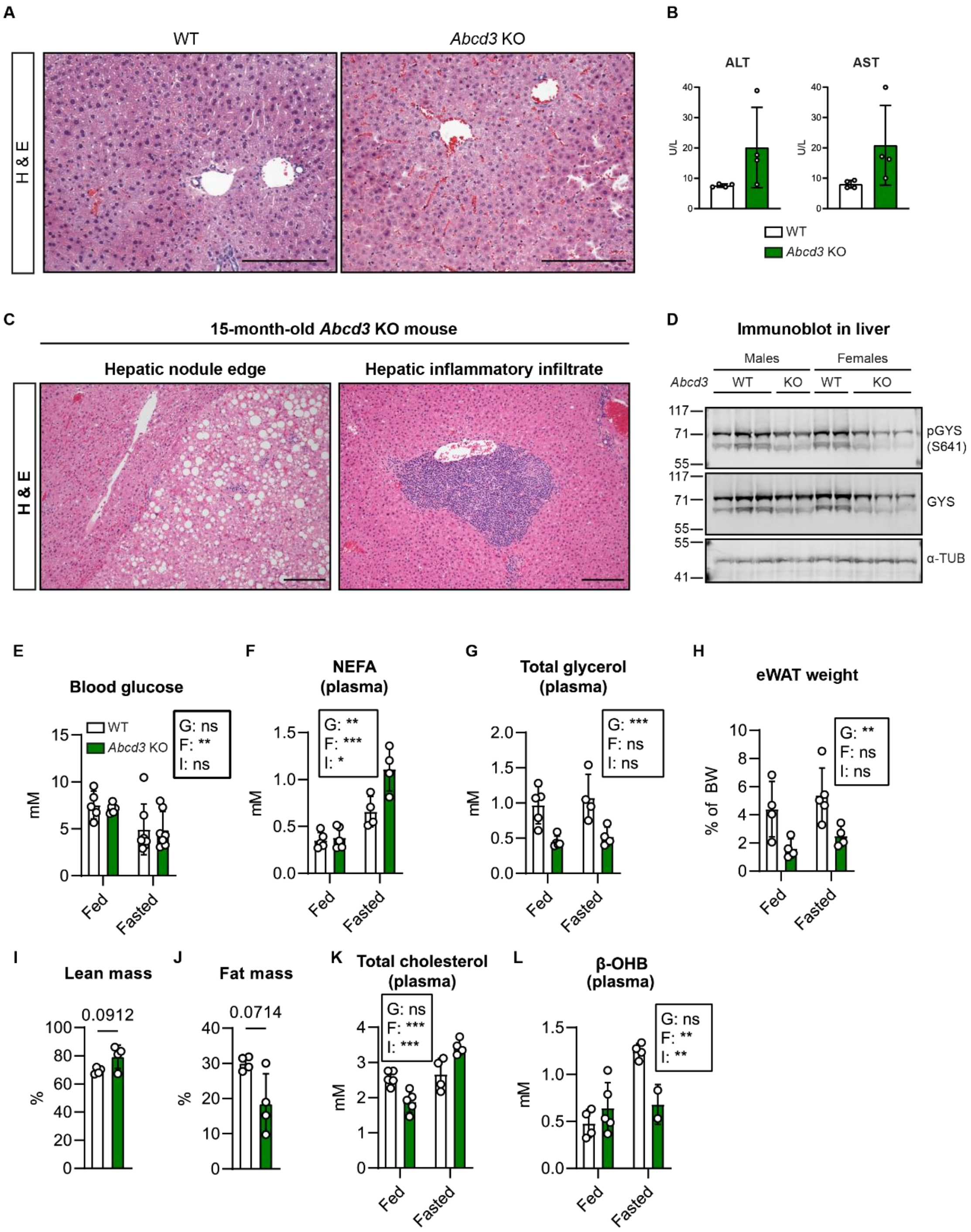

Supplementary Figure 2

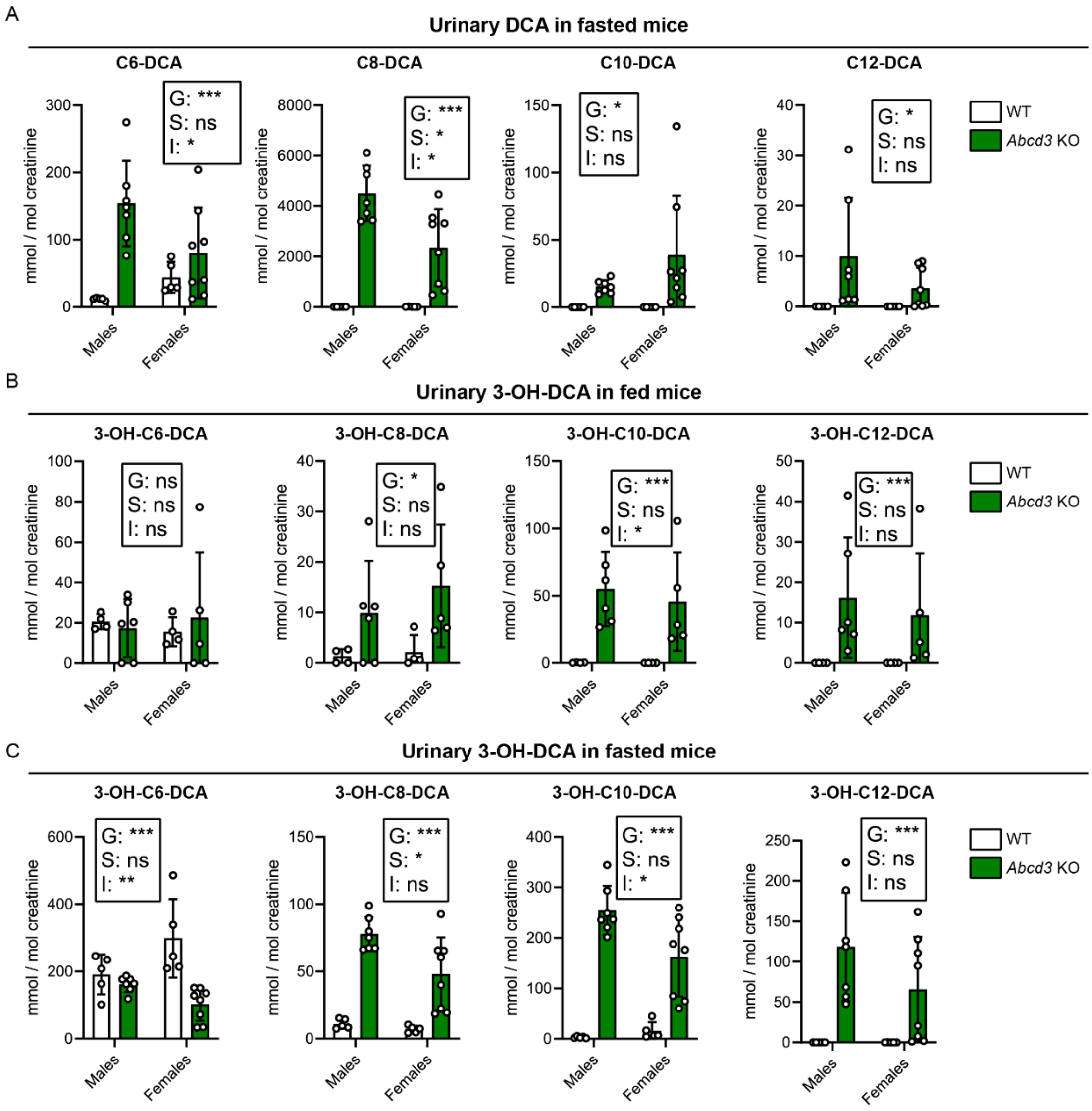

Supplementary Figure 3

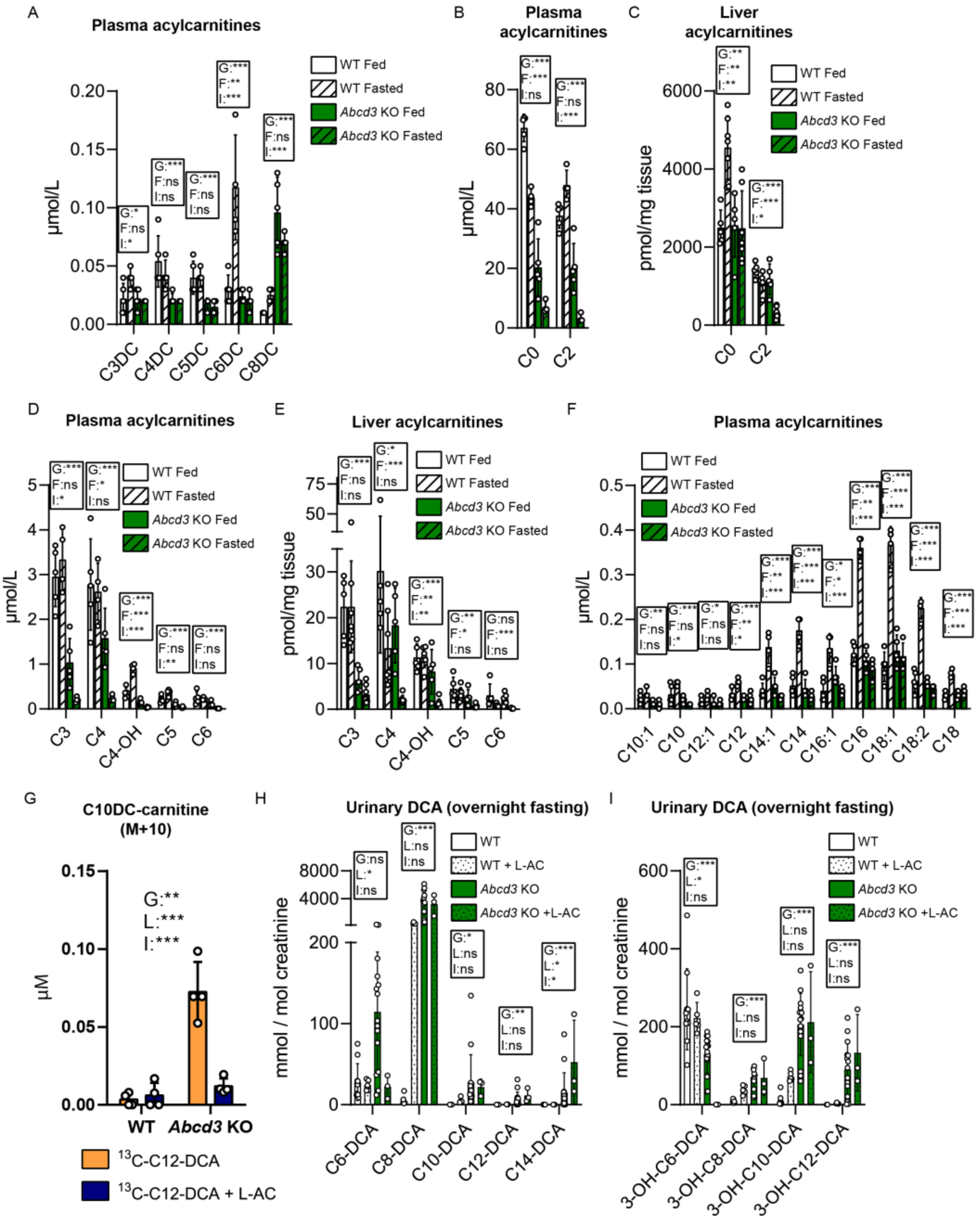

*Supplementary Figure Legends*

**Figure S1**. Characterization of a second *Abcd3* KO mouse

**A**) Representative images of H & E staining of WT and *Abcd3* KO livers from male fasted mice. Scale bar = 100 µm. **B**) Alanine aminotransferase (ALT) and aspartate aminotransferase (AST) activities (in U/L) in WT and *Abcd3* KO plasma (n=4). **C**) Representative images of H & E staining of an *Abcd3* KO liver that presented nodules with vesicular steatosis and inflammatory infiltrates. Scale bar = 100 µm. **E**) Blood glucose levels (in mM). **F**) Non-esterified fatty acids (NEFA) levels (in mM) in plasma. **G**) Total glycerol levels (in mM) in plasma. **H**) Epididymal white adipose tissue (eWAT) pad mass (as the % of body weight). BW: body weight. **I**) Lean mass (in %) as measured by echoMRI. **J**) Fat mass (in %) as measured by echoMRI. **K**) Total cholesterol levels (in mM) in plasma. **L**) Beta-hydroxybutyrate (β-OHB) levels (in mM) in plasma. Individual values, the average and the standard deviation are graphed for each parameter.* $p<0.05$; ** $p<0.01$; *** $p<0.001$ (unpaired, two-tailed student t-test in B, I, J; two-way ANOVA in E-H, and K-L). G: Genotype; F: Feeding; I: Interaction.

**Figure S2**. Urinary DCA alterations in *Abcd3* KO mice

**A**) Urinary C6-DCA, C8-DCA, C10-DCA, and C12-DCA (in mmol/mol creatinine) in fasted WT (n=5 males, n=5 females) and *Abcd3* KO (n=7 males, n=8 females) mice. **B**) Urinary 3-OH DCAs (C6-, C8-, C10-, and C12-, in mmol/mol creatinine) in WT (n=4 males, n=4 females) and *Abcd3* KO (n=6 males, n=5 females) fed mice. **C**) Urinary 3-OH DCAs (C6-, C8-, C10-, and C12-, in mmol/mol creatinine) in WT (n=5 males, n=5 females) and *Abcd3* KO (n=7 males, n=8 females) fasted mice. Individual values, the average and the standard deviation are graphed for each parameter.* $p<0.05$; ** $p<0.01$; *** $p<0.001$ (two-way ANOVA). The effects in the two-way ANOVA are abbreviated as follows: G: Genotype; S: Sex; I: Interaction.

**Figure S3**. DCA metabolism alterations in *Abcd3* KO mice

**A**) Plasma short- and medium-chain dicarboxylylcarnitine profile (in µmol/L) in fed and fasted WT and *Abcd3* KO mice (n=5 per genotype and feeding condition). **B**) Plasma free carnitine (C0) and acetylcarnitine (C2) levels (in µmol/L) in fed and fasted WT and *Abcd3* KO mice (n=5 per genotype and feeding condition). **C**) Liver free carnitine (C0) and acetylcarnitine (C2) levels (in pmol/mg tissue) in WT (n=5 fed, n=7 fasted) and *Abcd3* KO (n=5 fed, n=7 fasted) mice. **D**) Plasma short-chain acylcarnitine profile (in µmol/L) in fed and fasted WT and *Abcd3* KO mice (n=5 per genotype and feeding condition). **E**) Liver short-chain acylcarnitine profile (in pmol/mg tissue) in WT (n=5 fed, n=7 fasted) and *Abcd3* KO (n=5 fed, n=7 fasted) mice. **F**) Plasma medium- and long-chain acylcarnitine profile (in µmol/L) in fed and fasted WT and *Abcd3* KO mice (n=5 per genotype and feeding condition). **G**) Measured [U-$^{13}$C]-labeled C10-DC-carnitine (in µmol/L) in media of mouse liver slices after 4-hr incubation of WT and *Abcd3* KO mouse liver slices (n=4) with [U-$^{13}$C]-C12-DCA alone or with [U-$^{13}$C]-C12-DCA + L-aminocarnitine (L-AC). **H**) Urinary even-chain DCAs (C6- to C14-, in mmol/mol creatinine) in WT (n=10 vehicle, n=5 L-AC) and *Abcd3* KO (n=15 vehicle, n=3 L-AC) mice injected with vehicle or L-aminocarnitine and subjected to overnight food withdrawal. **I**) Urinary even-chain 3-OH-DCAs (C6- to C12-, in mmol/mol creatinine) in WT (n=10 vehicle, n=5 L-AC) and *Abcd3* KO (n=15 vehicle, n=3 L-AC) mice injected with vehicle or L-AC and subjected to overnight food withdrawal. Individual values, the average and the standard deviation are graphed.* p<0.05; ** p<0.01; *** p<0.001 (two-way ANOVA). The effects in the two-way ANOVA are abbreviated as follows: G: Genotype; F: Feeding; L: L-AC; I: Interaction.
